# Nutraceutical supplement targeting multiple molecular steps synergistically enhances muscle glucose uptake and improves in vivo oral glucose disposal

**DOI:** 10.1101/2022.07.09.499288

**Authors:** Uday Saxena, Kranti Meher, Saranya K, Arpitha Reddy, Gopi Kadiyala, Subramanian Iyer, Subrahmanyam Vangala, Satish Chandran

## Abstract

Pre diabetes and type 2 diabetes are increasingly becoming rampant world wide. While there are medications to control blood glucose in type 2 diabetics, currently there are no interventions prescribed for pre diabetes. Alternate strategies to control blood glucose are needed either to act alone in pre diabetes or as supplement to the existing drugs for type 2 diabetes. We report here the targeting of critical molecular steps in muscle glucose uptake and metabolism to result in glucose lowering using a combination of safe vitamins. Our in vitro and in vivo data support the potential for using such vitamin combination for glucose control in pre diabetes and as a supplement in type 2 diabetics.

## INTRODUCTION

Type 2 Diabetes is a condition that impairs the body’s ability to process blood glucose. High glucose blood levels are responsible for the morbidities and life-style changes associated with diabetes. Type 2 diabetes is a lifelong disease with severe consequences if blood glucose is left uncontrolled. It may lead to irreversible kidney damage (nephropathy), nerve damage (neuropathy), damage to eyes (retinopathy), heart disease and stroke.

Prior to the onset of full-fledged type 2 diabetes, patients undergo a pre-diabetic phase which lasts for years during which there is a year-on-year increase in fasting blood glucose, eventually culminating in full-fledged diabetes if left unchecked. Figure 1 below shows the typical progression from normal blood glucose level to full fledged type 2 diabetes.

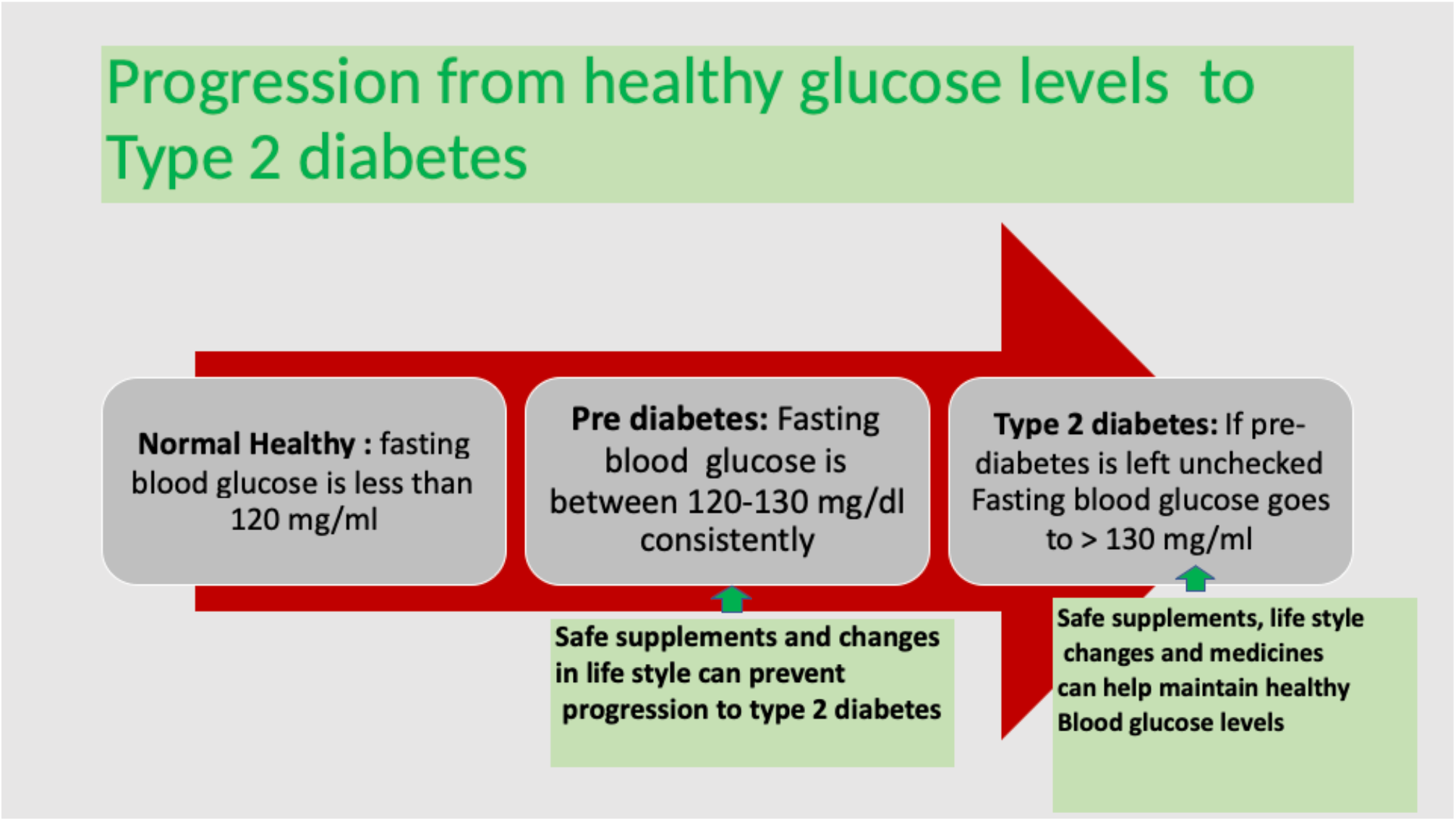

Type 2 diabetes is caused by diet and life style changes such as eating high calorie food, lack of exercise, obesity, family history and high stress being the common causes. In addition high carbohydrate foods lead to high blood sugar. Upon consumption of food, pancreas releases insulin which then triggers muscle taking up glucose and lowering blood sugar. Furthermore, age, stress, sedentary life style, over-eating makes the muscle not responding to insulin (insulin resistance) which leads to high blood glucose and eventually type 2 diabetes (1,2).

Although there are several medications available to treat type 2 diabetes, they all have limitations and multiple drugs have to be used for for maximum control. There is still a need for a safe nutraceutical supplement combination which can act at multiple molecular steps to help insulin action in muscle cells. To this end we have developed a mix of safe and well tolerated vitamins as nutraceutical supplement which act synergistically at various steps in the insulin signalling cascade and improve glucose uptake.

Normally, muscles take up almost 70% of the glucose from blood which leads to blood glucose lowering. The hormone insulin produced by the islet cells in pancreas is the main trigger for this glucose uptake by muscles. In pre-diabetes and type 2 diabetes, muscles become non-responsive to insulin (3,4). Any intervention that can help promote insulin induced muscle glucose uptake will help balance blood glucose levels (Figure 2 below shows glucose uptake pathway)

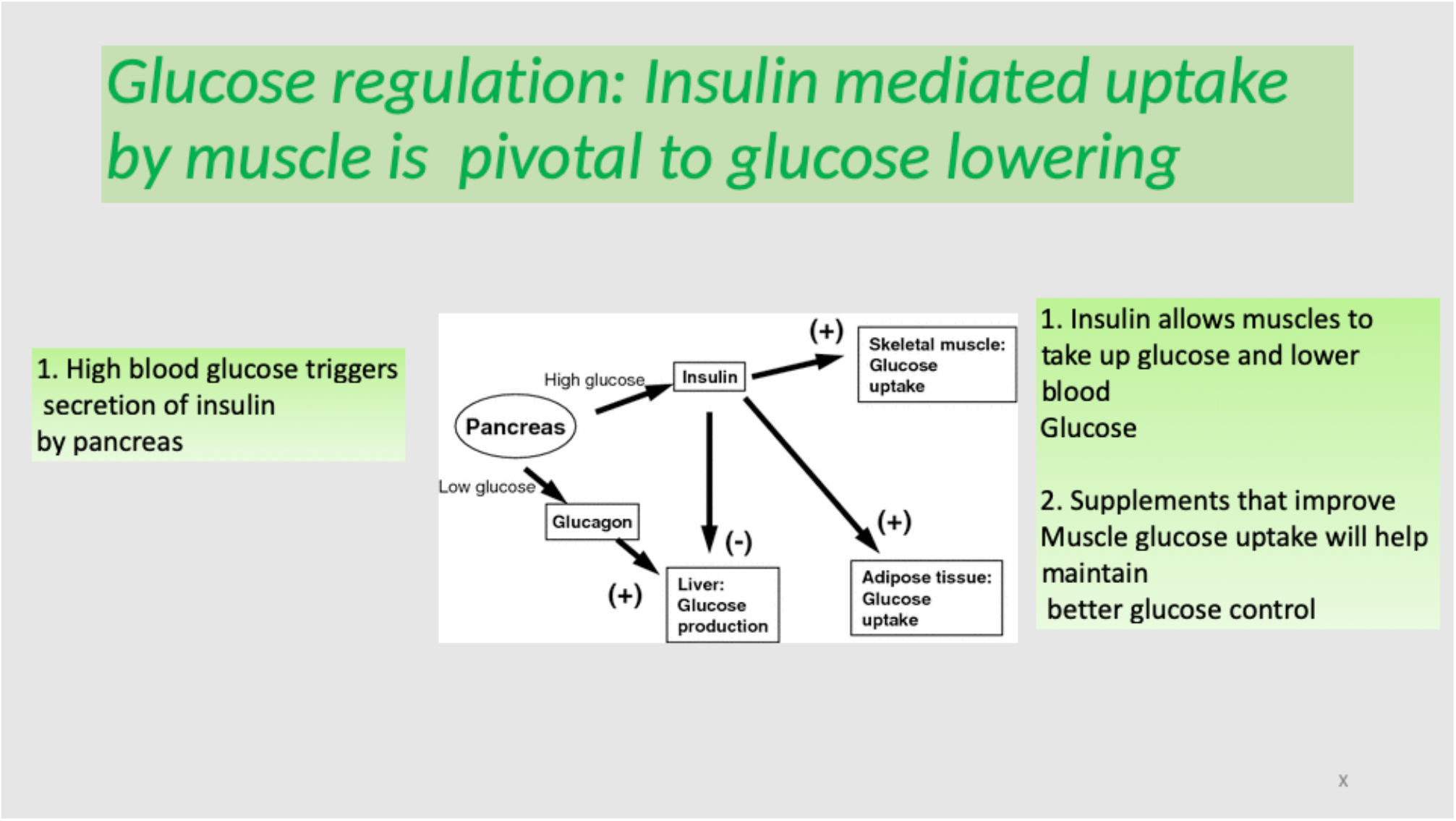

Once carbohydrate food is ingested, the dietary carbohydrates get broken down in the gut into glucose, which is then absorbed by the intestinal cells into the blood circulation. Within a short time of glucose absorption, the pancreas releases insulin triggering a cascade of steps that clear glucose for circulation. Essentially the steps are that 1) insulin binds to its receptors in various tissues such as muscle and adipose. 2) This then recruits the glucose transporters GLUT4 from cytoplasm to the cell membrane where it binds glucose and transports it inside the cells. 3) Once inside the cell’s glucose can either be utilised to generate energy (ATP) or be stored as glycogen. The uptake of glucose by the tissues lowers blood glucose levels (4,5). The various molecular steps in glucose uptake by muscle are:

1. Insulin triggers the uptake of glucose into cells by triggering relocation of GLUT4 transport protein
2. The glucose is then used in the mitochondria to generate energy thru a series of redox reactions which need nicotinamide
3. The redox reactions generate oxidative stress which if not controlled can inhibit ATP generation

Given the need for safe and effective interventions we explored the use of targeted vitamins which can improve muscle glucose uptake. We demonstrate here that a judicious mix of these in a fixed dose combination markedly enhance glucose uptake in an in vitro model and also blunts the rise in plasma glucose in an oral glucose tolerance test in rats.

## Methods

Glucose uptake studies using L6 cells were performed similar to our previous reports (6). The concentrations of Vitamin D3, Niacinamide and Lipoic acid were typically similar to the the Required Daily Allowance (RDA) for each. Briefly L6 cells (ATCC CLR-1458; Rockville, MD) were maintained in Dulbecco’s modified Eagle’s medium (DMEM) supplemented with 10% Fetal Bovine Serum (FBS) and 100 μ g/mL penicillin and streptomycin, at 37 °C in an atmosphere of 5% CO2.2-NBDG glucose uptake. 2-(N-[7-nitrobenz-2-oxa-1, 3-diazol-4-yl] amino)-2-deoxyglucose (2-NBDG) (Molecular Probes-Invitrogen, CA, USA) was used to assess glucose uptake in L6 cells. Cells were kept in glucose free medium for half an hour before insulin stimulation. Cells were stimulated with 100 nM insulin for 10 and 20 min and incubated with 10 μ M of 2-NBDG for 15 min. The reaction was stopped by washing with cold PBS three times and the cells were lysed 0.1% Triton X. The lysate was then used to read florescence at 535 nM. The data is reported as percent of control cells (no treatment, incubation with media).

Oral Glucose tolerance test in male wistaria rats was performed similar to what we have reported previously (7). The studies were performed by Liveon Biolabs PVT Ltd under IAEC approval. The protocol used briefly was as follows : animals were acclimatized and were randomly distributed into 6 groups, each group comprised of 6 rats. The rats in group G1 serves as normal control group and administered with Vehicle alone at the dose volume of 10 mL/kg whereas the rats in G4 group were administered with Vitamin mix respectively at the doses of 200 mg/kg body weight for Single administration.respectively at the dose of half capsule a day for two weeks. All the Rats were observed for clinical signs, morbidity, mortality, body weight and blood glucose were measured during the study period.

### Test System Details

|  |  |
| --- | --- |
| Animal Species | : Rat (*Rattus norvegicus*) |
| Strain | : Wistar Rats |
| Justification for Selection of Species | : Rats are used as a standard laboratory rodent species for pharmacological studies as per literature and research articles. |
| Source | : In-house bred animals |
| Age at treatment | : 7 – 8 weeks |
| Total number of animals | : 12 |
| Sex | : Male |
| Number of animals per group | : 6 |
| Number of groups | : 2 |
| Bodyweight range ( at treatment) | : Around 250 grams |

### Animal Room Preparation

Prior to housing the animals, the experimental room was decontaminated by fumigation and microbial load was checked by settle plate method. The experimental room floor was mopped with disinfectant daily once.

### Husbandry Conditions

Animals were housed in an environment-controlled room at temperature of 20.1 - 22.5 °C and relative humidity of 52 - 68%. The photoperiod was 12 hours fluorescent light and 12 hours darkness. Adequate fresh air supply of 12 - 15 air changes/hour and sound level of <80 dB was maintained in the experimental room.

The relative humidity, Maximum and minimum temperature in the experimental room were recorded once daily. The copies of results is included in the study file.

### Housing

Two rats per cage were housed in standard polycarbonate cage (size: approximately Length 43 x Breadth 29 x Height 18 cm) with stainless steel top grill having facilities for pelleted food and drinking water. During the study period, rats were housed in a single experimental room. Polycarbonate rat tunnels were provided to the animals as an environmental enrichment objects in the cages that either provide shelter or exercising opportunities to minimize animal stress and promote overall well-being. These enrichment objects were replaced at least once a week.

#### Test Procedure

##### Wistar Rats were Randomly Divided into 2 Groups

For two weeks vehicle (Ethanol + water) and test item were administered for respective groups. After 30 min of last dose, glucose were administered in the dose 2gm/kg. 0min (before glucose administration), after glucose administration 30min, 60min, 90min, 120min and 180min was measured glucose level by using Accu Chek glucometer.OGTT study was conducted on overnight fasting animals.The dose formulation preparation were prepared freshly before administration and were used within 2 hours of formulation preparation.

##### 2 week repeated dose OGTT study

Vitamin mixture consisting of RDA levels of Vitamin D3, Niacinamide and lipoic acid dissolved in 2 ml of of ethanol (50%) and 1 ml of water and half of the volume used to dose the animals orally for two weeks.

##### Diet and Water

AF-1000M R&M diets manufactured by Krishna Valley Agrotech LLP was provided ad libitum to rats.

Deep bore-well water subjected to reverse osmosis and UV sterilized, was provided ad libitum to rats in polycarbonate bottles with stainless rubber corked steel sipper tubes. The food and water provided to the Rats were tested for contaminants. Acclimatisation

The animals were acclimatized to experimental room conditions for a period of 6 days prior to initiation of treatment. Body weights were recorded at the initiation and end of acclimatization period. Animals were observed for mortality and morbidity once daily during acclimatization period. Veterinary examination was performed before selecting the animals and only healthy and active animals were used in the study.

##### Animal Identification

During acclimatization period (Temporary identification), each animal was identified by animal number written on tail using marker pen. The cages were identified with cage cards indicating study number, study code, species, strain, sex, acclimatization start and acclimatization end date.

During treatment period (Permanent identification), each animal was identified by tail marking with animal accession number written on tail using permanent marker pen. The cages were identified with cage cards indicating study number, animal accession number, study code, species, strain, sex, treatment start date and treatment end date.

##### Randomization and Grouping

Grouping of animals was performed on the fifth day of acclimatization by body weight randomization and stratification method and allocated into two groups G1 and G4 groups, The body weight variation within the groups of animals did not exceed ±20% of the mean body weight. Body weights of the animals were analyzed statistically for mean body weight to rule out the statistically significant difference between groups.

### In VIVO STUDY COMPLIANCE

- The study was performed as per the mutually agreed Study Plan and the Standard Operating Procedures (SOPs) of the Test Facility.
- IAEC APPROVAL

The use of animals for this study had been approved by Liveon Biolabs Private Limited Institutional Animal Ethics Committee (IAEC). IAEC approved protocol number LBPL-IAEC-014-01/2021.

Animal welfare and veterinary care

Liveon Biolabs Private Limited is an AAALAC International accredited facility and registered with CPCSEA, Department of Animal Husbandry and Dairying (DAHD), Ministry of Fisheries, Animal Husbandry and Dairying (MoFAH&D), Government of India. Also, Liveon Biolabs Private Limited ensures that animal experiments are performed in accordance with the recommendation of the regulatory guidelines for laboratory animal facility published in the gazette of India, 2021.

During the conduct of study none of the animals were injured, and no moribund animals were observed.

## RESULTS

### 1. Effect of Vitamin D3 and Niacinamide on glucose uptake by muscle cells

Our goal was to improve insulin’s ability to increase the uptake of glucose by L6 muscle cells. We first targeted two specific molecular events in glucose uptake 1) increasing GLUT4 levels in the cells to enhance glucose transport into the cells by insulin and 2) use up the glucose for energy generation in the mitochondria where using TCA cycle glucose is metabolised to generate ATP by using niacinamide (a precursor of nicotinamide adenine dinucleotide, NAD a co-enzyme in TCA cycle). Replenishment of NAD can further boost glucose uptake and and utilisation.

We used Vitamin D3 (8,9) to increase GLUT4 levels since this vitamin has been shown before to increase GLUT4 levels in muscle and used added Niacinamide to increase cellular NAD levels (10,11).

As shown in Figure 3 we found that in presence of insulin in control cells, not treated with VitaminD3 or Niacinamide, there was only modest uptake of glucose at 10 minutes and none at 20 minutes after insulin pulse treatment.

**Figure 3:**
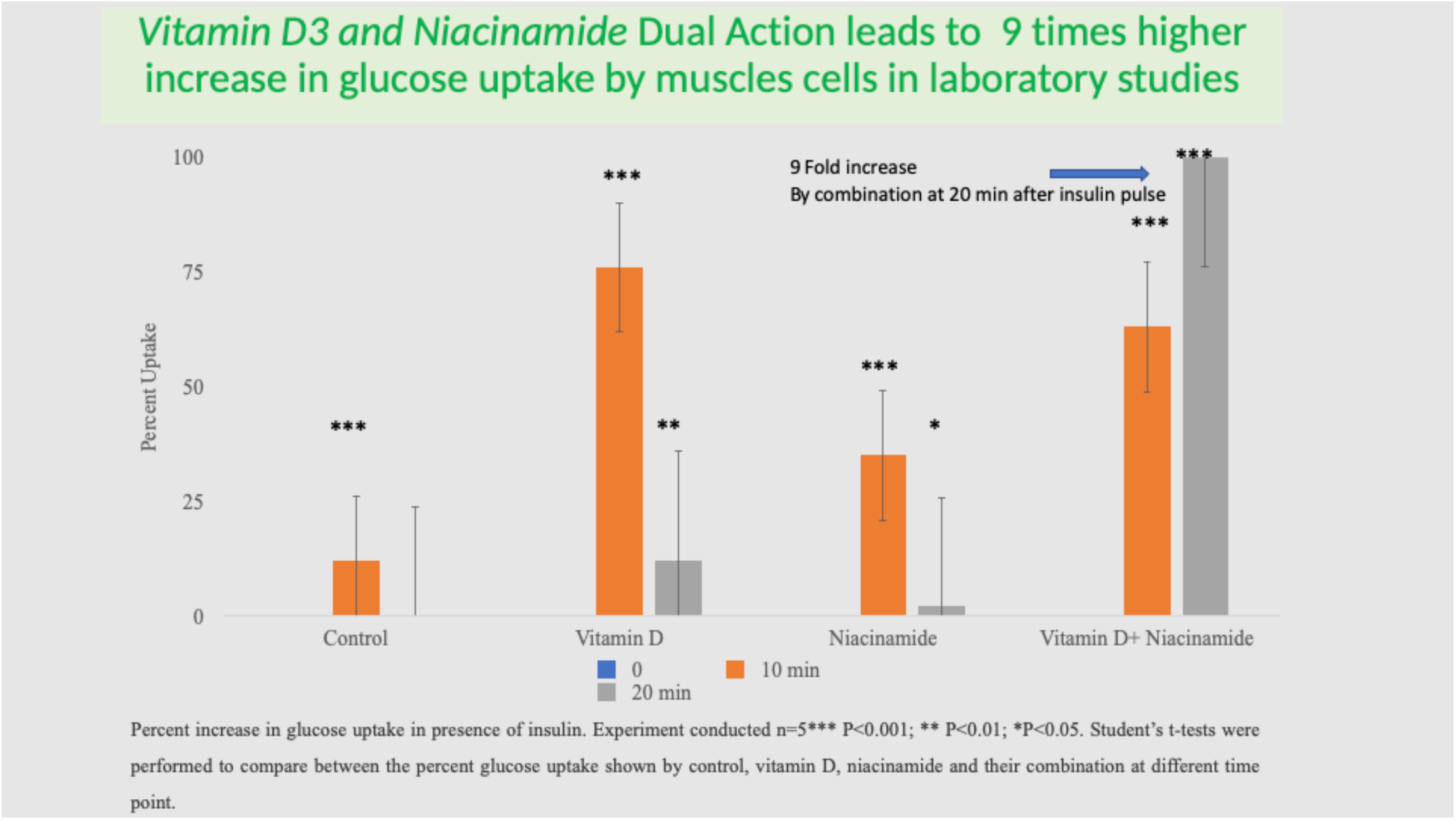

When Vitamin D3 alone was added, it increased glucose uptake after insulin pulse by about 6-fold more than control cells at 10 minutes but less so at 20 minutes. Thus suggest that Vitamin D3 alone can increase glucose uptake.

We then tested the effect of niacinamide treatment. Niacinamide increased glucose uptake by about 3 fold over control at 10 minutes but only little at 20 min, suggesting that this vitamin can also increase glucose uptake on its own.

We then combined the two agents to see if there is a synergistic effect since they work thru different mechanisms. Figure 3 shows a very dramatic increase in glucose uptake by the combination especially at 20 minutes where the increase was 9 fold higher and more that either of the agents suggesting synergy. The other interesting thing is that the combination statins the glucose increase peaks 20 minutes unlike when either agent is added alone.

These data suggest a synergistic effect go Vitamin D and Niacinamide in glucose uptake by muscle cells.

### 2. Effect of adding lipoic acid

Since the TCA cycle and glucose metabolism in the mitochondria generate oxidative stress, we next explored if lipoic acid, an antioxidant (12) which is known to enter the mitochondria can further enhance glucose uptake.

We tested the following combinations for improvements in muscle cell glucose uptake stimulated by insulin:

- Vitamin D3 plus Niacinamide plus or minus Lipoic acid alone

As shown in Figure 4. Above relative to control cells and as expected from above studies the combination of VitaminD3 and niacinamide (V=N) showed significant increase in glucose uptake relative to control cells. In contrast Lipoic acid alone did not increase glucose uptake alone but the combination of Vitamin D3, Niacinamide and Lipoic acid (LA+V+N) increased glucose uptake even moderately better than Vitamin D3 and Niacinamide combination. Interestingly Vitamin E a powerful antioxidant but which does not enter the mitochondria had no effect. These data support the idea of improvement in glucose metabolism by suppressing mitochondrial oxidative stress.

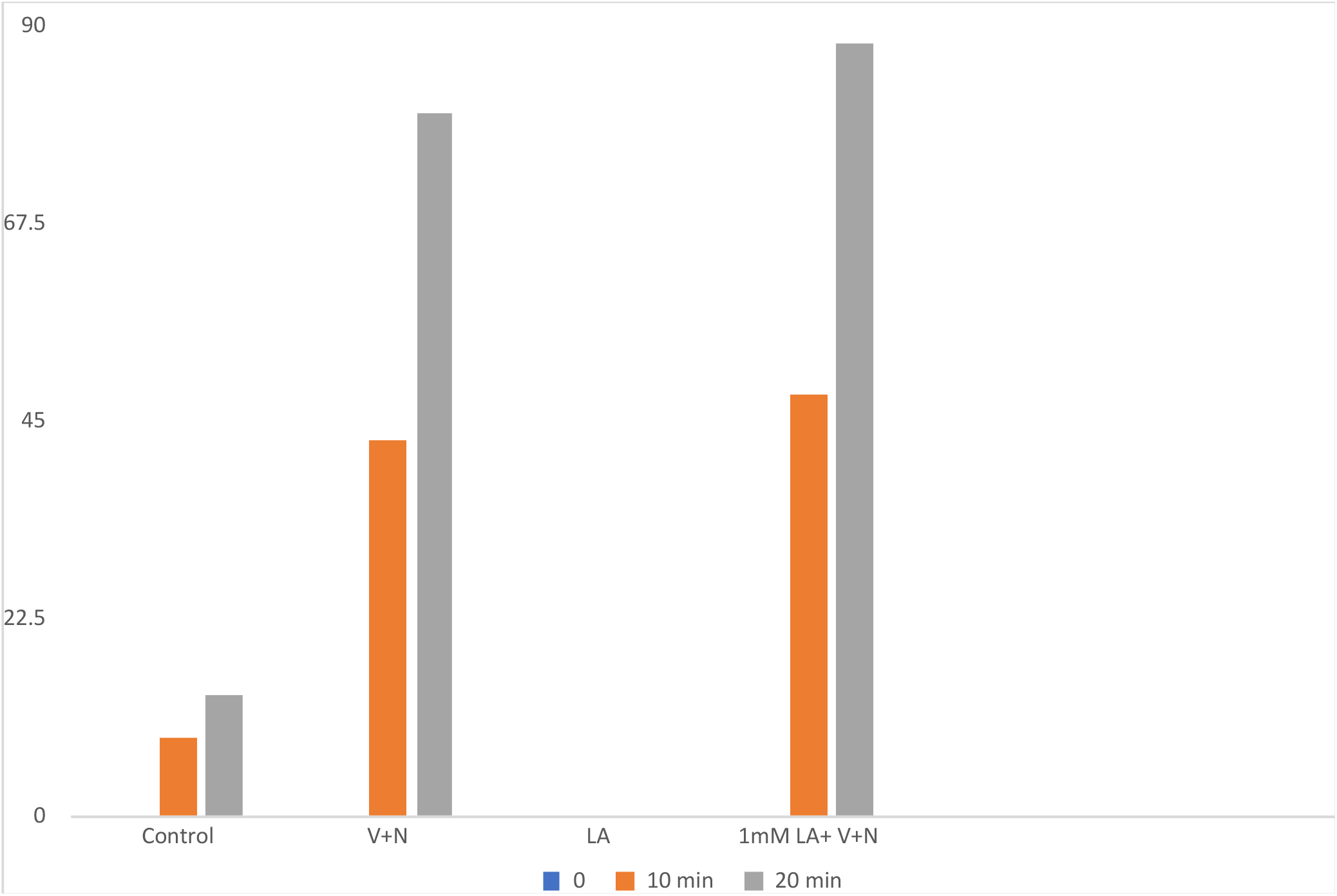
*The data shown above are statistically significant for the V+N and LA+V+N from control p>0.05*

### 3. Oral Glucose Tolerance Test (OGTT) in Vitamin mixture treated rats

Oral glucose tolerance test (OGTT) is often used to test the effect of an intervention on disposal of orally given glucose as a measure of the body’s ability to clear glucose from the circulation. The lower the glucose levels after a glucose meal, the more effective is the physiological system including insulin, muscle, liver and adipose etc in glucose uptake.

As shown below in figure 5 below after oral glucose load at every time point examined, there was statistically significant decrease in the vitamin treated group in glucose levels relative to the vehicle treated control group. These data suggest that the vitamin has improved the glucose disposal system from circulation and resulted in glucose lowering.

**Figure 5:**
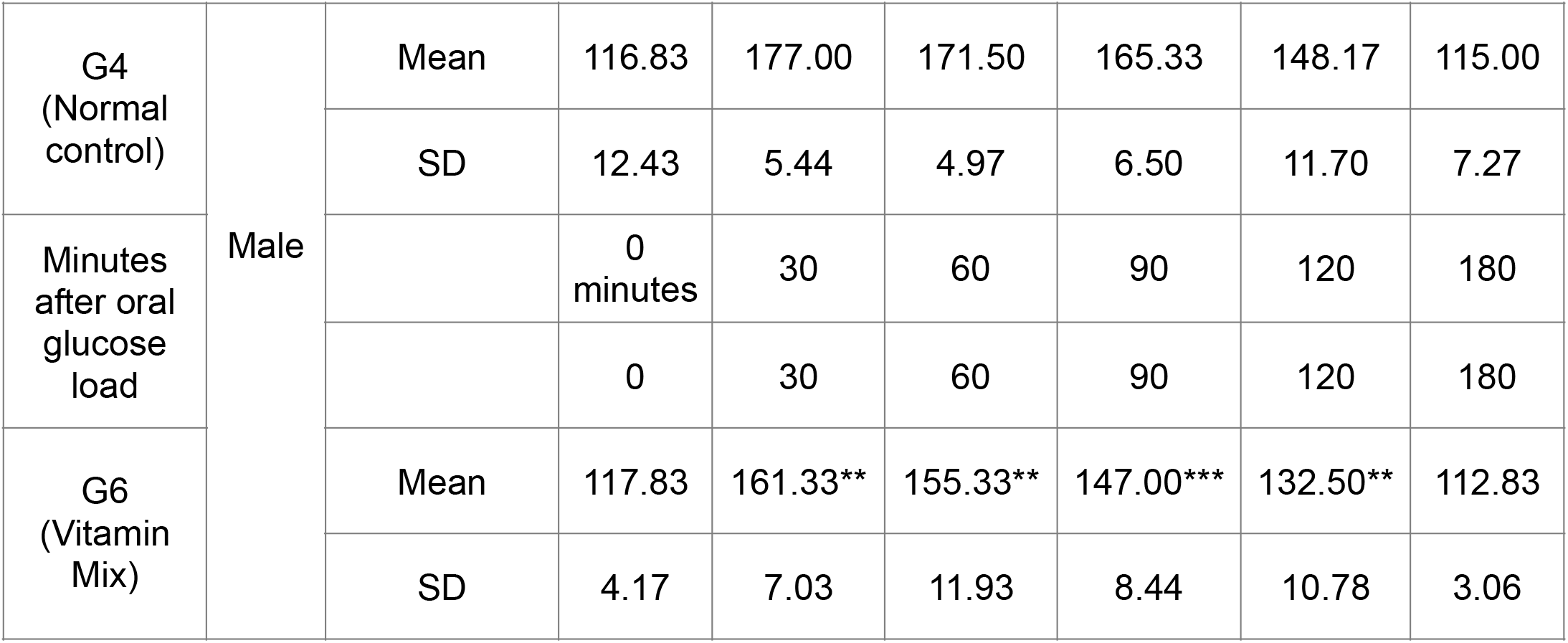
Summary of blood glucose levels in animals after Oral Glucose Tolerance Test Data expressed in Mean ± SD: (n=6) *P<0.01, **P<0.001, ***P<0.0001 VS control group

The calculated Area Under the Curve (ACU) of glucose levels from 0-180 minutes time point also showed a statistically significant decrease suggesting that the Vitamin mix effect was sustained and successfully blunted the rise in blood glucose after oral glucose load (Figure 6)

**Figure 6:**
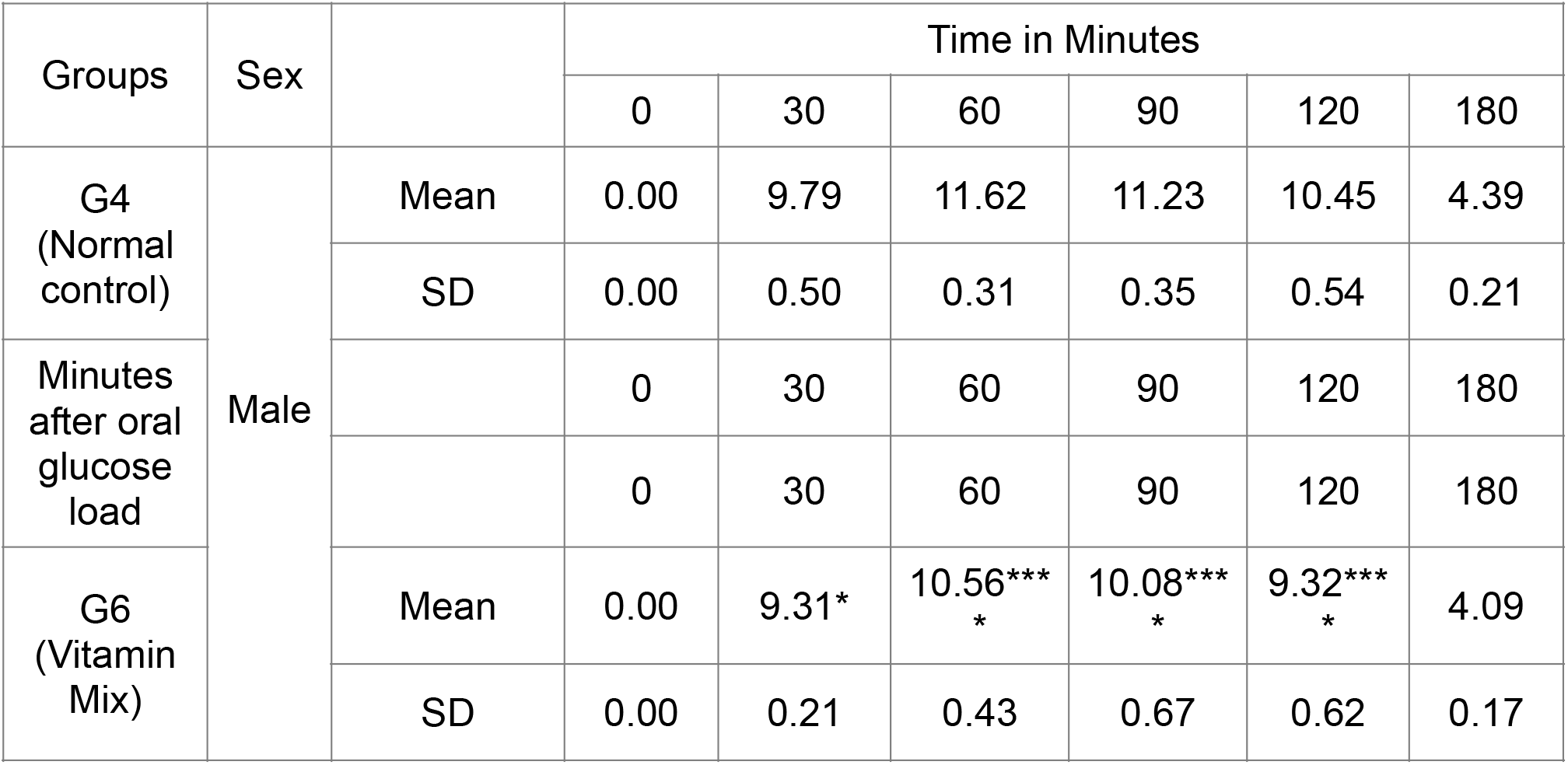
Summary of Area Under the Curve (AUC) values for blood glucose The values are expressed in Mean ± SD: (n=6) *P<0.01, **P<0.001, ***P<0.0001 as compared to induction control (G4) group.

## DISCUSSION

The current studies were initiated to identify interventions that had large well know safety data in humans and could intervene at rate limiting molecular events that improve glucose uptake and metabolism by tissues. Given the need for established safety we focused our attention on the Vitamins that have used for decades or longer.

To this end we identified glucose uptake and metabolism in cells as important drivers of blood glucose lowering. Specifically we focussed on GLUT4 transporter which imports glucose into the cells after insulin receptor binding.Vitamin D3 has been reported to increase GLUT4 levels in muscle cells and was our candidate molecule to test.

Secondly we proposed that the glucose which is transported in cells has to be metabolised completely and efficiently to produce glucose lowering effect. For this we explored niacinamide which boost cellular NAD levels a critical Co-enzyme for glucose metabolism via TCA cycle and lipoic acid an antioxidant which enters the mitochondria and is able to suppress any oxidative stress which may shut down glucose metabolism. To the best of our knowledge this is the first time that a rational use of vitamin combination in a specific ratio has been shown to effect glucose lowering. Our belief is that such alternative intervention will be of great value addition for glucose control in pre diabetics and as well as can be used as adjunct supplement in type 2 diabetics.

